# Three-dimensional experiments and individual based simulations show that cell proliferation drives melanoma nest formation in human skin tissue

**DOI:** 10.1101/244582

**Authors:** Parvathi Haridas, Alexander P. Browning, Jacqui A. McGovern, D.L. Sean McElwain, Matthew J. Simpson

## Abstract

**Background:** Melanoma can be diagnosed by identifying nests of cells on the skin surface. Understanding the processes that drive nest formation is important as these processes could be potential targets for new cancer drugs. Cell proliferation and cell migration are two potential mechanisms that could conceivably drive melanoma nest formation. However, it is unclear which one of these two putative mechanisms plays a dominant role in driving nest formation.

**Results:** We use a suite of three-dimensional (3D) experiments in human skin tissue and a parallel series of 3D individual-based simulations to explore whether cell migration or cell proliferation plays a dominant role in nest formation. In the experiments we measure nest formation in populations of irradiated (non-proliferative) and non-irradiated (proliferative) melanoma cells, cultured together with primary keratinocyte and fibroblast cells on a 3D experimental human skin model. Results show that nest size depends on initial cell number and is driven primarily by cell proliferation rather than cell migration.

**Conclusions:** We find that nest size depends on initial cell number, and is driven primarily by cell proliferation rather than cell migration. All experimental results are consistent with simulation data from a 3D individual based model (IBM) of cell migration and cell proliferation.

## Background

Clusters of melanoma cells, called *nests*, are a prominent feature of melanomas [1,2]. Identifying the presence and characteristics of melanoma nests in human skin is an essential diagnostic tool [3,4]. Recent 3D experimental work by Wessels *et al*. [5] suggests that melanoma nest formation in Matrigel is driven by cell migration. However, nest formation might be different in human skin, where melanoma cells are in contact with other cell types [1,6]. We hypothesise that two different mechanisms could lead to nest formation: (i) cell proliferation, where clusters of melanoma cells are formed primarily through mitosis (Figure 1A); and (ii) cell migration, where clusters of adhesive melanoma cells form primarily through melanoma cell migration (Figure 1B).

**Figure 1:**
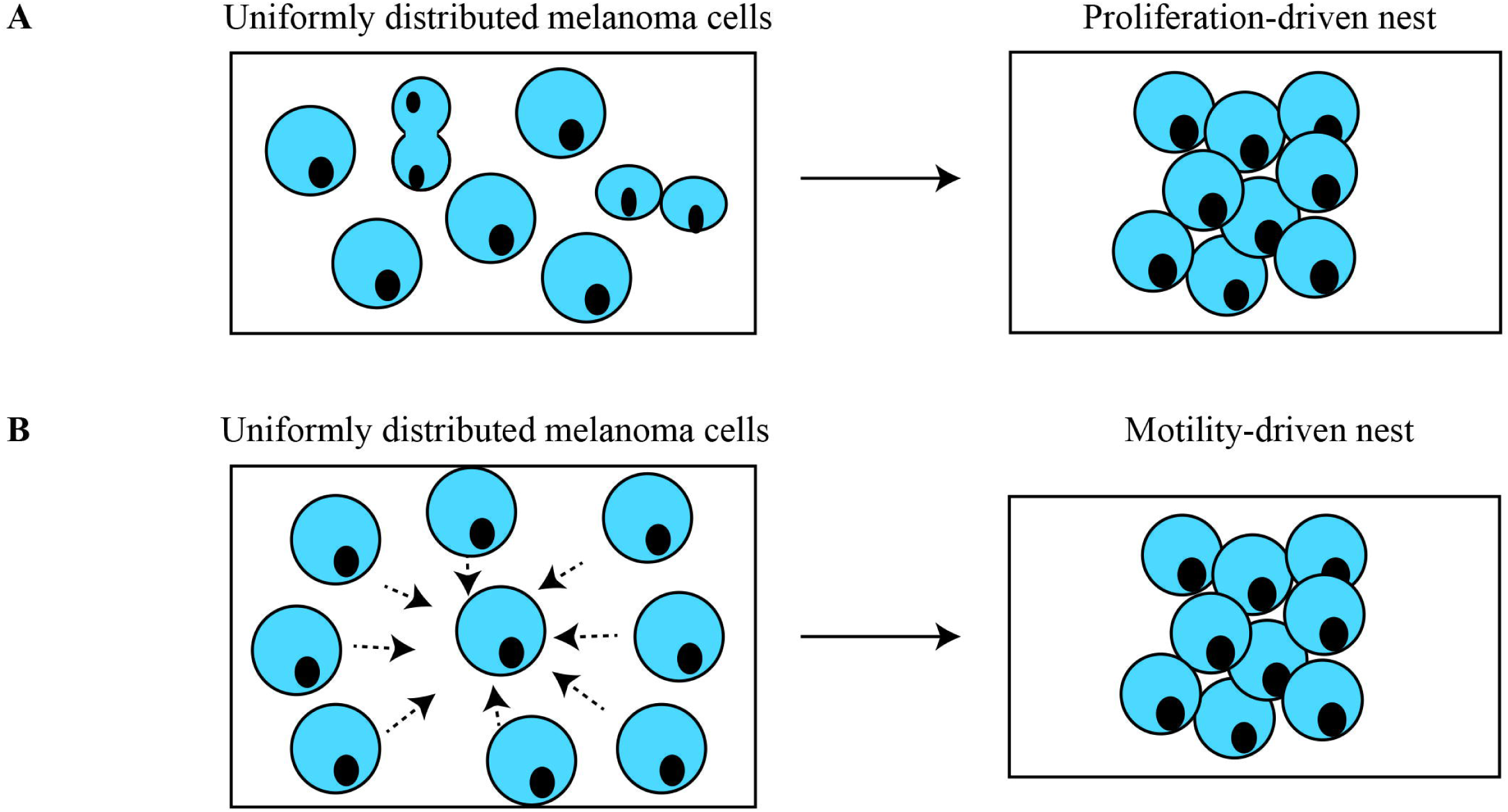
Mechanisms that drive melanoma nest formation. Schematics illustrating: (A) proliferation-driven nests; and (B) migration-driven nests. In both cases the schematic shows an initially-uniform distribution of cells that lead to the formation of a nest either by the action of proliferation (A) or migration (B).

We use a 3D human skin experimental model [7,8] to discriminate between these two conceptual models by performing a suite of experiments in which we systematically vary the initial density of proliferative melanoma cells placed on 3D human skin. This initial series of experiments allow us to examine the role of initial cell number in driving nest formation. All experiments are then repeated using non-proliferative, gamma-irradiated melanoma cells. We find that higher initial numbers of melanoma cells lead to larger nests, and that cell proliferation leads to dramatically-larger nests. All experimental outcomes are consistent with a series of 3D simulations from an IBM [9]. These results provide insight into the mechanisms driving nest formation, showing that the mechanisms in 3D human skin are different to monoculture experiments performed in Matrigel.

## Results and Discussion

### Confirmation that irradiated melanoma cells do not proliferate and are capable of migrating in a two-dimensional barrier assay

Experiments involving populations of proliferative melanoma cells are performed using non-irradiated SK-MEL-28 cells [10]. Experiments where melanoma cell proliferation is suppressed are performed using irradiated, but otherwise identical SK-MEL-28 cells [11,12]. The melanoma cells are gamma-irradiated to inhibit mitosis. We perform a series of live assays to show that irradiation does not affect the adherence or morphology of melanoma cells. Furthermore, the live assay confirms that irradiated melanoma cells do not proliferate, and that the irradiation does not cause the cells to die [see Additional file 1].

Two-dimensional (2D) barrier assays confirm that irradiated melanoma cells survive and migrate. Populations of irradiated melanoma cells are monitored over four days to confirm that irradiation does not impede the ability of cells to migrate. We use circular barrier assays to compare the spatial expansion of irradiated and non-irradiated melanoma cell populations. The leading edge of the spreading populations is detected using ImageJ [13], which also provides an estimate of the area occupied by the spreading population of cells. Since the spreading populations of cells maintain an approximately circular shape, we convert the estimates of area into an equivalent diameter and we report data in terms of the diameter of the spreading population. Results are obtained in triplicate. Images in Figure 2A-B show the increase in the diameter of the spreading cell populations for both irradiated and non-irradiated melanoma cells over four days. The upper row of images in Figure 2A-B, show increased spatial expansion of the population of non-irradiated cells compared to the population of irradiated melanoma cells in the lower row. Since irradiated melanoma cells do not proliferate, we expect that the size of the expanding population of irradiated cells will be smaller than the size of the expanding population of non-irradiated cells [14]. However, the area occupied by the population of irradiated melanoma cells increases over the four-day period, and this increase is due to cell migration alone. To confirm these visual observations, we also quantify the spatial spreading of irradiated and non-irradiated melanoma cell populations.

**Figure 2:**
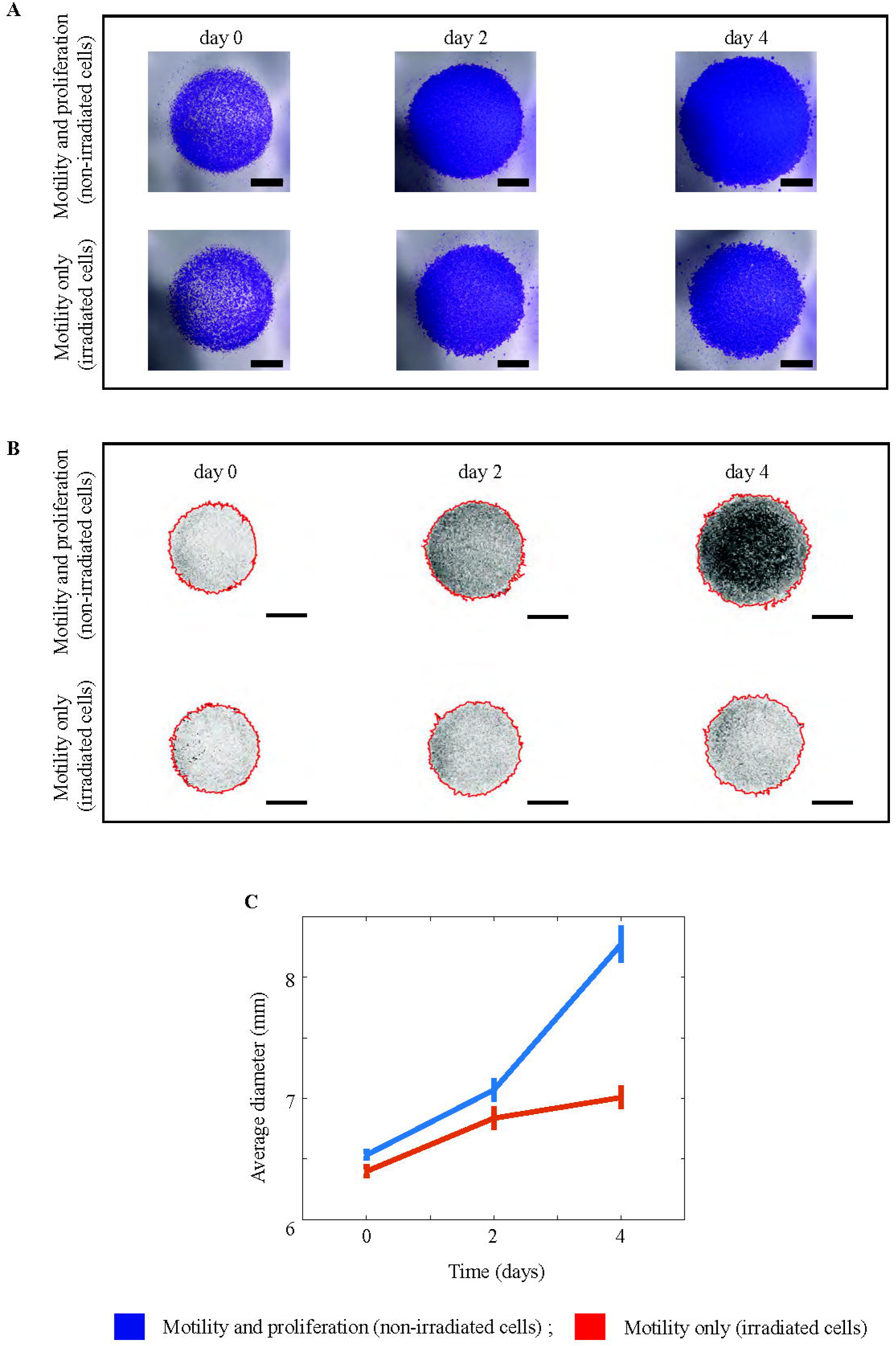
Two-dimensional spatial expansion of irradiated and non-irradiated melanoma cell monocultures. (A) Experimental images show barrier assays initialised with approximately 10000 melanoma cells. The upper row of images show non-irradiated (proliferative) melanoma cells, and the lower row shows irradiated (non-proliferative) melanoma cells. The images show the spreading of cell populations at zero, two and four days, respectively. The scale bar is 2 mm in each image. (B) Experimental images from (A) analysed by ImageJ. Results show the position of the leading edge of the spreading population (red) superimposed on images of the spreading populations. The upper row of images corresponds to non-irradiated melanoma cells, and the lower row of images show irradiated melanoma cells. The images show the spreading of cell populations at zero, two and four days, respectively. The scale bar in each image is 3 mm. (C) and (D) Data shows the average diameter of the spreading populations as a function of time (n=3). All data generated using non-irradiated melanoma cells is in blue, and data generated using irradiated melanoma cells is in red. Plots in (D) also show the variability. The error bars correspond to the sample standard deviation (n=3).

Data in Figure 2C shows the increase in diameter of both irradiated and non-irradiated melanoma cell populations. At all times considered, the average diameter of the irradiated cell population is less than the average diameter of the non-irradiated cell population. This is expected because the irradiated melanoma cells do not proliferate, and it is known that proliferative populations of cells expand and invade the surrounding empty space faster than non-proliferative populations of cells [9,14]. Most importantly, the experiments initialised with irradiated melanoma cells show an increase in the diameter of the spreading population, confirming that irradiation does not prevent migration. All experiments are performed in triplicate and the averaged results are presented. We now use both, irradiated and non-irradiated melanoma cells in 3D experiments to identify the mechanism that drives melanoma nest formation.

### Identifying the dominant mechanism driving melanoma nest formation

Nests of melanoma cells are well-characterised histological features of melanoma progression. Early identification of these nests is critical for successful melanoma treatment. However, in addition to examining the presence of melanoma nests, it is important to identify the biological mechanisms that lead to nest formation as this information might be relevant to the development of new drugs. To examine these pathways we use a 3D experimental skin model.

Irradiated and non-irradiated melanoma cells are cultured with primary keratinocytes and primary fibroblasts in the 3D experimental skin model for four days. From this point we refer to keratinocyte and fibroblast cells as *skin* cells. All cells are initially placed onto the 3D experimental skin model as a monolayer, as uniformly as possible. MTT (3-(4,5-dimethylthiazol-2-yl)-2,5-diphenyltetrazolium bromide) assays highlight the metabolic activity of all cells, and show the spatial extent and spatial structure of cells on the top surface of the 3D experimental skin model. Images in Figure 3A-B show prominent dark purple clusters on the surface of some 3D experimental skin models. Control studies, where 3D experiments are constructed without melanoma cells, show a complete absence of nests [see Additional file 1] suggesting that the dark purple clusters in Figure 3A-B are melanoma nests.

**Figure 3:**
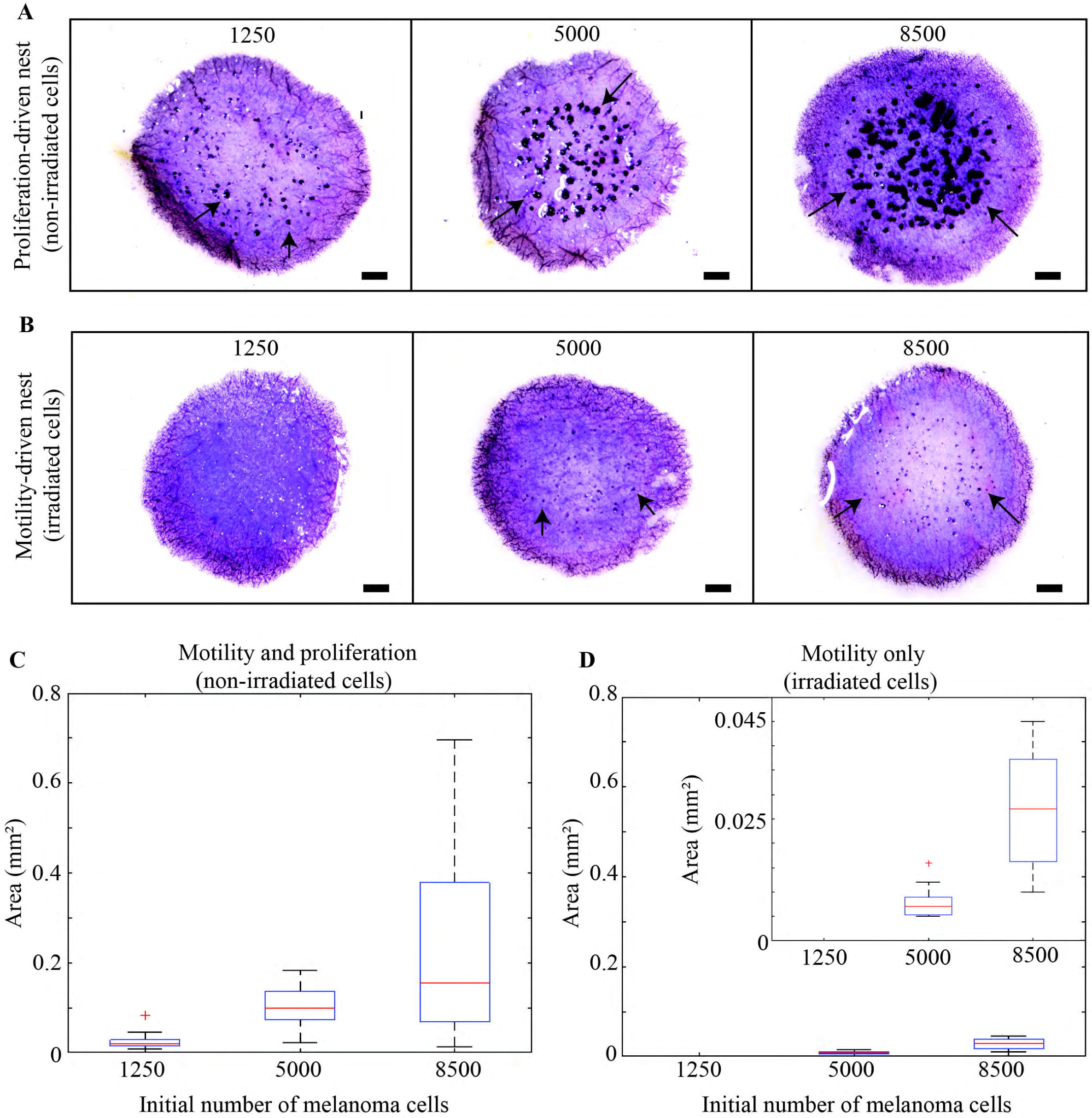
Proliferation drives melanoma nest formation. (A) MTT assays show all metabolically active cells (light purple) on the surface of the 3D experimental skin model initialised with different numbers of proliferating melanoma cells, as indicated. (B) Equivalent results with irradiated melanoma cells. Melanoma nests are in dark purple (arrows). Scale bars are 1 mm. (C)-(D) Box plots showing nest area as a function of initial number of melanoma cells. Inset in (D) shows details in the range 0-0.045 mm^2^.

Images in Figure 3A show that larger nests are associated with higher initial numbers of melanoma cells. To quantify this we measure the area of individual nests using ImageJ [13], and data in Figure 3C confirms our visual observation. Interestingly, larger initial numbers of melanoma cells lead to a smaller number of larger nests [see Additional file 2]. This is consistent with smaller sized nests coalescing into a smaller number of larger nests over time. Since cell number plays a critical role, we will also examine the role of proliferation by suppressing mitosis.

We examine the role of cell proliferation by constructing 3D experimental skin models with irradiated melanoma cells. Images in Figure 3B show that this leads to the formation of dramatically smaller nests. To quantify our results, the area of individual nests is measured using ImageJ [13] [see Additional file 2]. Data in Figure 3D shows a similar trend to data in Figure 3C as the nest area increases with initial cell number. However, comparing results in Figure 3C-D shows that proliferation plays a dominant role in nest formation. For example, experiments initialised with 8500 proliferative melanoma cells leads to a median nest area of 0.15 mm^2^, whereas the median nest area is just 0.027 mm^2^ when proliferation is suppressed.

Our results are different to previous 3D studies that show melanoma nests are formed by cell migration [5]. We anticipate that the difference in our outcome could be due to: (i) differences between the melanoma cell lines used; (ii) the interaction of melanoma cells with the surrounding skin cells in our 3D experiments; or, (iii) differences in the material used to construct the 3D model. Since our experiments are performed in 3D materials derived from human skin, and our experiments involve culturing melanoma cells together with primary human skin cells, we feel that our results are more realistic than examining nest formation in monoculture experiments in Matrigel. We now perform immunohistochemistry to confirm that irradiated melanoma cells in survive in the 3D experimental human skin model over a period of four days.

### Irradiated melanoma cells survive in a 3D experimental skin model

Here, we perform a series of experiments using a specific melanoma marker to provide additional evidence that nests observed on the 3D experimental human skin models are clusters of melanoma cells, and that irradiated melanoma cells survive in a 3D environment over four days. The 3D experimental skin models are constructed using both irradiated and non-irradiated melanoma cells. Vertical cross-sections through the 3D experimental skin models initialised with melanoma cells are stained using S100, which is a reliable melanoma cell marker [15]. Both irradiated and non-irradiated melanoma cells are found in the 3D experimental skin model after four days. Images in Figure 4A-F show positive S100 staining of melanoma cells. In particular, Figure 4B,D,F show positive S100 staining of irradiated melanoma cells after four days. This immunostaining confirms that irradiation does not alter the antigen properties of melanoma cells, and the irradiated melanoma cells survive in a 3D experimental skin model for four days. Our experimental results use skin cells and skin dermis from one donor. Additional results using cells and dermis from two other donors show little variability between them.

**Figure 4:**
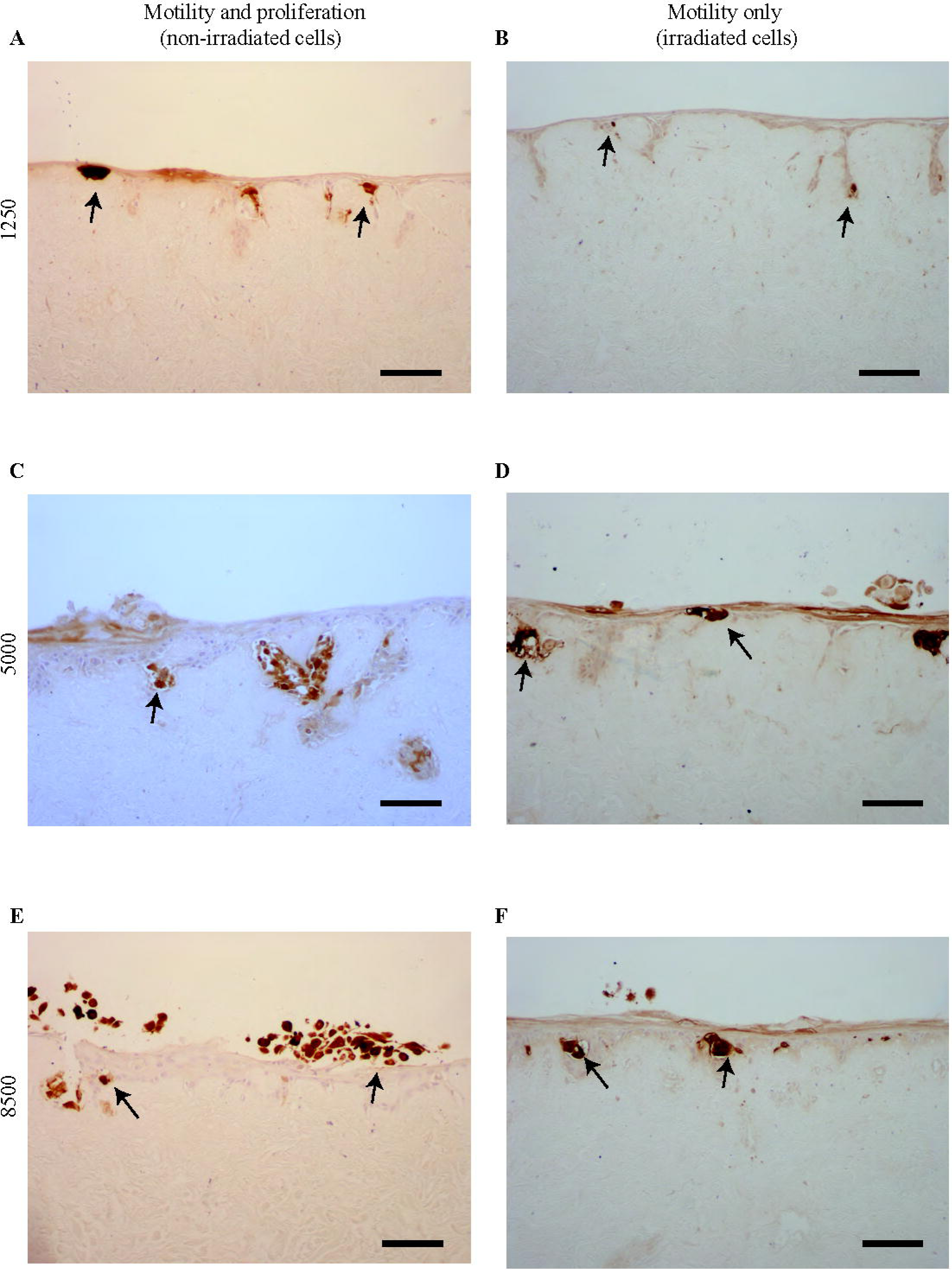
Irradiated and non-irradiated melanoma cells survive in 3D experimental skin models. S100 identifies melanoma cells (brown), and the arrows indicate positive staining. (A), (C) and (E) Cross-sections through 3D experimental skin models initialised with 1250, 5000 and 8500 non-irradiated melanoma cells, as indicated. (B), (D), and (F) Cross-sections through 3D experimental skin models initialised with 1250, 5000 and 8500 irradiated melanoma cells, respectively. Scale bar in each image is 100 μm.

### Variability between skin samples

We now examine whether there is any important variability in our results between skin samples from different donors. To examine this we perform additional experiments using dermis and primary skin cells from three different donors, which we denote as donor A, donor B and donor C. We show MTT assays on the 3D experimental skin models initialised with non-irradiated and irradiated melanoma cells in Figure 5. The upper row of images in Figure 5A-C show 3D experimental skin models initialised with 1250, 5000 and 8500 non-irradiated melanoma cells, respectively. In each case, we see that larger nests are associated with higher initial number of melanoma cells. A similar trend is observed for the images in the lower row of images in Figure 5A-C where the experiments are initialised with an equivalent number of irradiated melanoma cells. However, regardless of whether we consider results from donor A, donor B or donor C, we always see that nest formation is dramatically reduced when we consider irradiated, non-proliferative melanoma cells.

**Figure 5:**
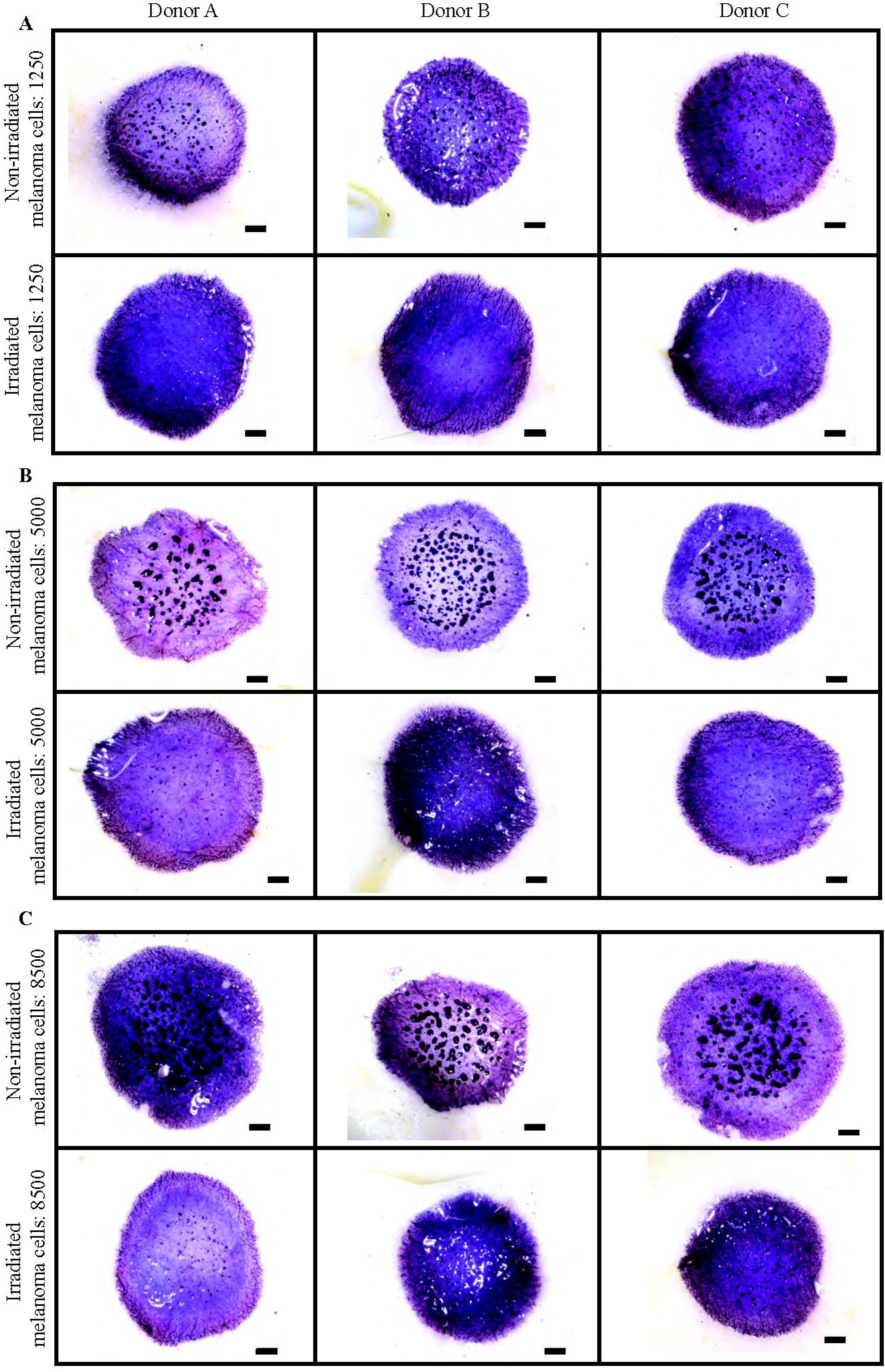
Donor variability in 3D experimental skin models with melanoma cells. Experimental images show metabolically active cells (light purple) on the 3D experimental skin model after four days. The skin models are constructed using primary skin cells and dermis from three different donor samples denoted A; B; and C. The scale bars are 1 mm. The melanoma nests are shown in dark purple. In each set of subfigures, (A)-(C), the images in the upper row show experiments initialised with 1250, 5000 and 8500 non-irradiated melanoma cells, respectively. In each set of subfigures, (A)-(C), the images in the lower row show experiments initialised with 1250, 5000 and 8500 irradiated melanoma cells, respectively.

Visual inspection of the images in Figure 5 suggests that the size, shape and number of individual nests does vary slightly between the three donors. However, the influence of the initial cell number and the action of cell proliferation on nest formation remains consistent between the skin samples from the three different donors. That is, larger initial numbers of cells produces larger nests, and the action of cell proliferation leads to dramatically larger nests. To provide additional evidence we also measure the area of individual nests on skin samples from all donors using ImageJ [13]. Data provided [see Additional file 2] confirm that the relationship between initial cell number and the action of cell proliferation holds for all three donor samples.

The nests on the 3D experimental skin model initialised with 1250 irradiated melanoma cells are very small. Most experimental replicates of this particular experiment do not lead to any visually observable nests, as shown in the lower row of images in Figure 5A. Therefore, data for nest area in these experiments is omitted [see Additional file 2]. We now use an IBM to verify our experimental outcomes.

### Modelling melanoma nest formation using an individual based model

To corroborate our experimental findings, we use an IBM to simulate the key features of the experiments. It is well-known that it can be difficult to quantitatively calibrate stochastic IBMs to match complicated experimental data precisely [16]. Therefore we use parameters in the IBM that are adapted from previous work [9,17]. This approach allows us to focus on understanding the roles of the key underlying biological features, such as the role of cell migration and cell proliferation, without being distracted by the secondary task of obtaining precise parameter estimates. We achieve this by using previously determined parameter estimates and simply comparing simulation results where melanoma cell proliferation is present, with simulation results where melanoma cell proliferation is suppressed.

The IBM describes the spatial distribution of simulated cells on a 3D square lattice [18]. We use a 3D lattice of cross section 3 mm × 3 mm, and depth 2 mm, to represent the central region of each experimental 3D skin model (Figure 6A). Simulated cells are called *agents*. We consider non-adhesive skin agents (green, Figure 6B) and adhesive melanoma agents (blue, Figure 6B).

**Figure 6:**
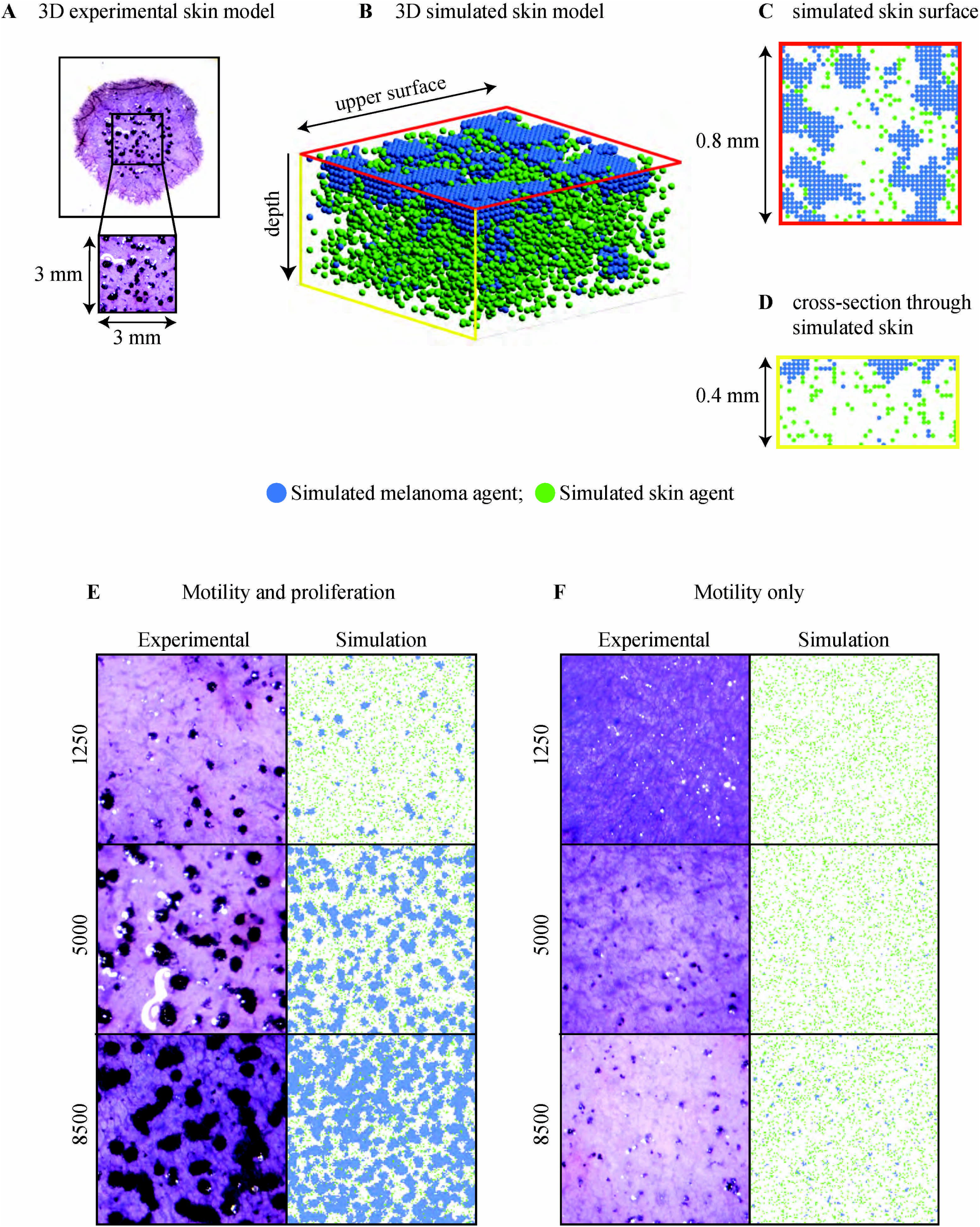
IBM simulations corroborate experiments. (A) Experimental image showing all metabolically active cells (light purple) on a 3D experimental skin model initialised with 5000 proliferating melanoma cells. The magnified 3 mm × 3 mm region shows melanoma nests (dark purple). (B) Sub-region of the 3D simulated skin model with simulated skin agents (green) and simulated melanoma agents (blue). The dimension of the upper surface is 0.8 mm × 0.8 mm, and the depth is 0.4 mm. (C) Upper surface of the simulated skin model. (D) Cross-section through the simulated skin model. (E)-(F) Experimental and simulated nests initiated with varying numbers of melanoma cells, as indicated, and an equivalent density of simulated melanoma agents, respectively. Results in (E) correspond to non-irradiated (proliferative) melanoma cells/agents. Results in (F) correspond to irradiated (non-proliferative) melanoma cells/agents. Images in (E)-(F) have dimensions 3 mm × 3 mm, and the depth is 2 mm. IBM parameters are *τ* = 0.01 h; Δ = 20 µm; *P_p_*^(^*^m^*^)^ = 0.0004; *P_m_*^(^*^m^*^)^ = 0.0075; *P_p_*^(^*^s^*^)^ = 0.00025; *P_m_*^(^*^s^*^)^ = 0.0075; and *q* = 0.7. Simulations with suppressed melanoma proliferation use *P_p_*^(^*^m^*^)^ = 0.0.

To initialise the IBM simulations, we randomly place a particular number of skin and melanoma agents onto the surface of the 3D lattice. The initial number of agents in each subpopulation is chosen to match to the initial cell density in the experiments. Figures 6B-D show smaller sub-regions of the 3D simulated skin to visualise the distribution of agents on the 3D lattice as clearly as possible. Results in Figure 6B-C show that the IBM predicts the formation of clusters of adhesive melanoma agents on the surface of the 3D lattice. Results in Figure 6D shows how the IBM predicts the downward movement of both skin and melanoma agents. Results in Figure 6D show that skin agents move deeper into the 3D lattice than the melanoma agents, while nests of melanoma agents tend to remain on the surface. Overall, the spatial arrangement of skin and melanoma agents in the IBM (Figure 6B-D) is similar to the spatial arrangement of cells in the 3D experiments (Figure 3–4) [6].

To explore the role of initial melanoma cell number in nest formation, IBM results in Figure 6E show that nests form on the surface of the 3D lattice, and that the trends in simulated nest area are similar to those in the corresponding experiments. Therefore, the simulation outcomes in Figure 6E confirm that initial melanoma cell number is an important factor in driving nest formation. We also explore the role of cell proliferation by repeating the simulations in Figure 6E without any melanoma agent proliferation. Simulation results in Figure 6F are comparable to the corresponding experimental results, as we observe similar trends in nest size and morphology. In conclusion, similar to the experiments, our 3D simulation results indicate that melanoma nest formation is driven by initial melanoma cell number, and that the presence of melanoma proliferation leads to dramatically-larger nests.

## Conclusion

Our combined experimental and simulation findings demonstrate that cell proliferation plays the dominant role in melanoma nest formation. While it is well-accepted that proliferation is important in the latter stages of tumour growth [19] and in the spatial spreading of cell populations [20], our work shows that proliferation is vitally important at the very earliest stages of melanoma progression.

Our results, pointing to the importance of cell proliferation, are interesting for a number of reasons: (i) previous monoculture experiments report that melanoma nests are formed by cell migration in Matrigel [5]. One potential explanation for this difference is that the Matrigel experiments are very different to our experiments since we study nest formation on 3D human tissues where melanoma cells are in contact with skin cells; (ii) some previous mathematical models of cluster/nest formation focus on cell migration only, e.g. [21], whereas we find that cell proliferation plays the most important role; and (iii) our findings about the importance of cell proliferation in melanoma progression are consistent with the fact that many promising melanoma drugs aim to suppress proliferation [22,23,24].

## Methods

### Keratinocyte isolation and culture

Queensland University of Technology (QUT) human research ethics obtained written approval for the skin samples to be used in this study (approval number: QUT HREC #1300000063; UnitingCare Health 2003/46). Skin samples are collected from patients undergoing elective plastic surgery. Human keratinocyte cells are isolated from skin and cultured in full Green’s medium following protocols described previously [15,25]. Primary keratinocyte cells are cultured at 37 °C, in 5% CO_2_ and 95% air.

### Fibroblast isolation and culture

Human fibroblast cells are isolated following protocols in Haridas *et al*. [15]. Primary fibroblast cells are cultured at 37 °C, in 5% CO_2_ and 95% air.

### Melanoma cell culture

The human melanoma cell line SK-MEL-28 is cultured as described in Haridas *et al*. [15]. SK-MEL-28 melanoma cells are kindly donated by Professor Brian Gabrielli (Mater Research Institute-University of Queensland). Cells are cultured at 37 °C, in 5% CO_2_ and 95% air.

A batch of SK-MEL-28 melanoma cells is irradiated to prevent cell proliferation. Approximately 1x10^7^ melanoma cells are gamma-irradiated using a Gammacell 40 research irradiator (Australia) at approximately 0.8 Gy/minute for one hour resulting in a cumulative dose of 50 Gy. We refer to these non-proliferative cells as *irradiated* melanoma cells, and the proliferative cells as *non-irradiated* melanoma cells.

Identification of SK-MEL-28 cells is validated using short tandem repeat profiling (Cell Bank, Australia. January 2015).

### Barrier assay

We perform circular barrier assays to observe and measure the spreading of populations of irradiated and non-irradiated melanoma cells. The protocol from Simpson *et al*. [14] is followed. Briefly, sterile stainless steel silicon barriers (Aix Scientific, Germany) are carefully placed in a 24-well tissue culture plate with 0.5 ml growth medium. The tissue culture plate containing cells is incubated for one hour at 37 °C, in 5% CO_2_ and 95% air. Viable cell suspensions of 20000 cells/100 µl of irradiated and non-irradiated melanoma cells are carefully introduced into the barriers to ensure an even distribution of cells. The tissue culture plates containing cell suspensions are incubated for a further two hours to allow cells to attach to the plate. The barriers are removed and the cell layers are washed with serum-free medium (culture medium without foetal calf serum) and replaced with fresh growth medium. Plates are then incubated at 37 °C, in 5% CO_2_ and 95% air for zero, two and four days. We replace the growth medium after two days to replenish the nutrients. Each assay is performed in triplicate.

### Crystal violet staining

We use the staining technique described by Simpson *et al*. [14] to analyse the barrier assays. In brief, cell monolayers are washed with phosphate buffered saline (PBS; Thermo Scientific, Australia) and fixed using 10% neutral buffered saline (United Biosciences, Australia) for 20 minutes at room temperature. The fixed cells are stained using 0.01% v/v crystal violet (Sigma Aldrich, Australia) in PBS for 20 minutes at room temperature. Excess crystal violet stain is removed using PBS, and the plates are air-dried. Images of irradiated and non-irradiated cell populations are acquired using a Nikon SMZ 800 stereo microscope fitted with a Nikon digital camera.

### Establishing 3D experimental skin model with melanoma cells

We establish 3D experimental skin models using the skin collected from donors undergoing elective plastic surgery. The protocol for establishing the 3D skin equivalent model with melanoma cells is adapted from previous work [7]. In brief, sterile stainless steel rings (Aix Scientifics) with a radius of 3 mm are placed on the papillary side of the de-epidermised dermis in a 24-well tissue culture plate (Nunc®, Australia). We refer to the de-epidermised dermis as *dermis*. Single cell suspensions of primary keratinocyte cells (20000), primary fibroblast cells (10000) and non-irradiated melanoma cells (1250; 5000; 8500), are seeded onto the dermis in full Green’s medium as uniformly as possible, and incubated at 37 °C, in 5% CO_2_ and 95% air for two days. We refer to the primary keratinocyte and fibroblast cells as *skin cells*. Subsequently, the stainless steel rings are removed and the dermis containing cells is submerged in full Green’s medium for a further two days. After this four-day pre-culture period, the spatial distribution of cells in the 3D experimental skin model is analysed. We also perform a series of equivalent experiments using irradiated melanoma cells.

All experiments are performed in triplicate. Furthermore, all experiments are repeated using primary skin cells and dermis from three separate donors to account for variability between different donors.

### MTT Assay

An MTT (Thermo Scientific) assay is performed to check the metabolic activity of cells on the 3D experimental skin models. These assays are imaged with a stereo microscope (Nikon SMZ 800) fitted with a Nikon digital camera. We follow the protocol from Haridas *et al*. [7].

### Immunohistochemistry on 3D experimental skin models with melanoma cells

We use immunohistochemistry to identify melanoma cells in the 3D experimental skin models. 10% neutral buffered formalin (United Biosciences, Australia) is used to fix the 3D experimental skin models. The tissue is divided through the centre of the MTT positive region using a sterile blade. The two smaller pieces of tissue are processed and embedded in paraffin. These samples are sectioned into 5 µm thick sections using a microtome. These sections are de-paraffinised, rehydrated and then subjected to heat-mediated antigen retrieval treatment using sodium citrate buffer (pH 6.0) in a decloaking chamber (Biocare Medical, USA) at 95 °C for 5 minutes. Skin sections are washed in PBS followed by immunostaining using the MACH 4™ Universal HRP polymer kit (Biocare Medical). The primary antibody S100 (Dako, Australia) is diluted in DaVinci Green diluent (Biocare Medical) at 1:3000, and these sections are incubated with the primary antibody for one hour at room temperature. Positive immunoreactivity is visualized using 3,3-diaminobenzidine (DAB; Biocare Medical) and then counterstained with using Gill’s haematoxylin (HD Scientific, Australia). The sections are dehydrated, and mounted on coverslips using Pertex® mounting medium (Medite, Germany). All stained sections are imaged using an Olympus BX41 microscope fitted with an Olympus digital camera (Micropublisher, 3.3RTV, QImaging; Olympus, Q-Imaging, Tokyo, Japan).

### IBM Simulation Methods

We use a 3D lattice-based IBM, with adhesion between some agents, to describe the 3D experiments. In the IBM, cells are treated as equally sized spheres, and referred to as *agents*. We use a square lattice, with no more than one agent per site. The lattice spacing, Δ, represents the approximate size of each simulated agent. Here, we set Δ = 20 µm. We use a 3D lattice of cross section, 3 mm × 3 mm, and depth 2 mm, to represent the central region of each experimental skin model. The parameters in the simulation model are adapted from previous studies [17]. Since we use the 3D lattice to represent the central region of the tissue, where cells are initialised uniformly across the surface, we apply periodic boundary conditions along all vertical boundaries. Since cells cannot leave the skin through the upper or lower surfaces, we apply no flux conditions on the upper and lower horizontal boundaries of the 3D lattice. We choose the depth of the 3D lattice to be large enough so that the agents never touch the bottom boundary of the lattice on the time scale of the simulations we consider.

To initialise simulations, we randomly place a particular number of simulated skin agents (*N*_0_^(^*^s^*^)^) and a particular number of simulated melanoma agents (*N*_0_^(^*^m^*^)^) onto the surface of the lattice. When the IBM is initialised we take care to ensure that no more than one agent occupies each lattice site. We always choose the initial number of agents in each subpopulation to match the equivalent initial density of cells in the experimental skin model. In the experiments, the initial populations of cells are uniformly placed inside a disc of radius 3 mm, whereas in the IBM the initial populations of agents are uniformly placed inside a square subregion of side length 3 mm. We set the initial number of skin agents to be *N*_0_^(^*^s^*^)^ = 9500 to match the initial experimental population of 30,000 skin cells distributed in a disc of radius 3 mm. We vary the initial number of simulated melanoma agents to be *N*_0_^(^*^m^*^)^ = 400, 1600 or 2700, to match the initial experimental populations of 1250, 5000 and 8500 melanoma cells distributed in a disc of radius 3 mm. To match the experiments, the IBM simulations run for four days.

At any time, *t*, there are *N*(*t*) agents on the lattice. In each discrete time step, of duration *τ*, we allow motility and proliferation events to occur in the following two sequential steps:

1. N(t) agents are selected one at a time, with replacement and given the opportunity to move to a nearest neighbour lattice site with probability *P_m_*^(^*^s,m^*^)^ ∈[0,1]. The probability of movement depends on whether the agent is a skin agent or a melanoma agent since we know that skin cells are more motile than melanoma cells [18]. If the chosen agent is a melanoma agent, we incorporate adhesion into the model by examining the occupancy of the 26 nearest lattice sites in the 3D von Neumann neighbourhood. We count the number of those sites occupied by melanoma agents, *a* [17]. Potentially motile melanoma agents then attempt to move with a modified probability, *P_m_^*^=* (1 − *q*)*^a^*, which accounts for adhesion between neighbouring melanoma agents. The parameter *q* controls the strength of melanoma-melanoma agent adhesion, with *q=0* corresponding to no adhesion, and increasing *q* leading to increased adhesion [18]. Setting *q=1* corresponds to maximal adhesion, and this would prevent any motility of melanoma agents that are in contact with other melanoma agents. We do not include any adhesion for the motion of skin agents. To simulate crowding effects, potential motility events that would place an agent on an occupied site are aborted.
2. *N*(*t*) agents are selected one at a time, with replacement and given the opportunity to proliferate with probability *P_p_*^(^*^s,m^*^)^ ∈[0,1]. The probability of proliferation depends on whether the agent is a melanoma agent or a skin agent [17]. If a proliferation event is successful, a daughter agent is placed at a randomly chosen nearest neighbour lattice site in the 3D Moore neighbourhood. To simulate crowding effects, we abort the proliferation event if all six nearest neighbouring sites are occupied. In all cases where a proliferation event is successful, a proliferative melanoma agent will produce a daughter melanoma agent, and a proliferative skin agent will produce a daughter skin agent.

The parameters in the IBM are Δ, *τ*, *P_m_*^(^*^s^*^)^, *P_m_*^(^*^m^*^)^, *P_p_*^(^*^i^*^)^ *P_p_*^(^*^m^*^)^ and *q*. These IBM parameters are related to the cell proliferation rates (*λ*^(^*^s^*^)^= *P_p_*^(^*^s^*^)^/ *τ*, λ^(^*^m^*^)^= *P_p_*^(^*^m^*^)^/ τ) and cell diffusivities (*D*^(*s*)^= *P_m_*^(^*^s^*^)^ Δ^2^/ (6*τ*), *D*^(^*^m^*^)^= *P_m_*^(^*^m^*^)^ Δ ^2^/(6τ)).

## Declarations

## Acknowledgements

We thank Brian Gabrielli for the SK-MEL-28 cell line.

## Availability of data and material

All data generated and/or analysed supporting the outcome of this manuscript is included within this manuscript and in the additional files.

## Authors’ contributions

PH and MJS conceived the study. PH, APB, JAM, DLSM and MJS designed the experiments. PH performed the experiments. APB performed the mathematical simulations. PH and APB analysed the data. PH and MJS wrote the manuscript. PH, APB, JAM, DLSM and MJS read, edited and approved the final manuscript.

## References

1. Beaumont KA, Mohana-Kumaran N, Haass NK. Modeling melanoma *in vitro* and *in vivo*. Healthcare. 2014;2:27–46. doi: 10.3390/healthcare2010027

2. Meier F, Nesbit M, Hsu M, Martin B, Belle PV, Elder DE, Schaumburg-Lever G, Garbe C, Walz TM et al. Human melanoma progression in skin reconstructs: biological significance of bFGF. Am J Pathol. 2000;156:193–200. doi: 10.1016/S0002-9440(10)64719-0

3. Balu M, Kelly KM, Zachary CB, Harris RM, Krasieva TB, Konig K, Durkin AJ, Tromberg BJ. Distinguishing between benign and malignant melanocytic nevi by in vivo multiphoton microscopy. Cancer Res. 2014;74:2688–2697. doi:10.1158/0008-5472.CAN-13-2582

4. Urso C, Rongioletti F, Innocenzi D, Batolo D, Chimenti S, Fanti PL, Filotico R, Gianotti R, Lentini M, Tomasini C, et al. Histological features used in the diagnosis of melanoma are frequently found in benign melanocytic naevi. J Clin Pathol. 2005;58:409–412. doi: 10.1136/jcp.2004.020933

5. Wessels D, Lusche DF, Voss E, Kuhl S, Buchele EC, Klemme MR, Russell KB, Ambrose J, Sol BA, Bossler A et al. Melanoma cells undergo aggressive coalescence in a 3D Matrigel model that is repressed by anti-CD44. PLoS ONE. 2017;12:e0173400. doi: 10.1371/journal.pone.0173400

6. Eves P, Layton C, Hedley S, Dawson RA, Wagner M, Morandini R, Ghanem G, Mac Neil S. Characterization of an *in vitro* model of human melanoma invasion based on reconstructed human skin. Brit J Dermatol. 2000;142:210–222. doi: 10.1046/j.1365-2133.2000.03287.x

7. Haridas P, McGovern JA, McElwain DLS, Simpson MJ. Quantitative comparison of the spreading and invasion of radial growth phase and metastatic melanoma cells in a three-dimensional human skin equivalent model. PeerJ. 2017;5:e3754. doi: 10.7717/peerj.3754

8. MacNeil S, Eves P, Richardson B, Molife R, Lorigan P, Wagner M, Layton C, Morandini R, Ghanem G. Oestrogenic steroids and melanoma cell interaction with adjacent skin cells influence invasion of melanoma cells *in vitro*. Pigment Cell Res. 2000;13:68–72. doi: 10.1034/j.1600-0749.13.s8.13.x

9. Treloar KK, Simpson MJ, Haridas P, Manton KJ, Leavesley DI, McElwain DLS, Baker RE. Multiple types of data are required to identify the mechanisms influencing the spatial expansion of melanoma cell colonies. BMC Syst Biol. 2013;7:137. doi: 10.1186/1752-0509-7-137

10. Carey TE, Takahashi T, Resnick LA, Oettgen HF, Old LJ. Cell surface antigens of human malignant melanoma: mixed hemadsorption assays for humoral immunity to cultured autologous melanoma cells. P Natl Acad Sci USA. 1976;73:3278–3282. doi: 10.1073/pnas.73.9.3278

11. Deacon DH, Hogan KT, Swanson EM, Chianese-Bullock KA, Denlinger CE, Czarkowski AR, Schrecengost RS, Patterson JW, Teague MW, Slingluff Jr CL. The use of gamma-irradiation and ultraviolet-irradiation in the preparation of human melanoma cells for use in autologous whole-cell vaccines. BMC Cancer 2008;8:360. doi: 10.1186/1471-2407-8-360

12. Todorovic D, Petrovic I, Todorovic M, Cuttone G, Ristic-Fira A. Early effects of gamma rays and protons on human melanoma cell viability and morphology. J Microsc-Oxford. 2008;232:517–521. doi: 10.1111/j.1365-2818.2008.02151.x

13. Schneider CA, Rasband WS, Eliceiri KW. NIH image to ImageJ: 25 years of image analysis. Nat Methods. 2017;9:671–675. doi:10.1038/nmeth.2089

14. Simpson MJ, Treloar KK, Binder BJ, Haridas P, Manton KJ, Leavesley DI, McElwain DLS, Baker RE. Quantifying the roles of cell motility and cell proliferation in a circular barrier assay. J R Soc Interface. 2013;10:20130007. doi: 10.1098/rsif.2013.0007

15. Haridas P, McGovern JA, Kashyap AS, McElwain DLS, Simpson MJ. Standard melanoma-associated markers do not identify the MM127 metastatic melanoma cell line. Sci Rep. 2016;6:24569. doi: 10.1038/srep24569

16. Read MN, Alden K, Rose LM, Timmis J. Automated multi-objective calibration of biological agent-based simulations. J R Soc Interface. 2016;13:20160543. doi: 10.1098/rsif.2016.0543

17. Haridas P, Penington CJ, McGovern JA, McElwain DLS, Simpson MJ. Quantifying rates of cell migration and cell proliferation in co-culture barrier assay reveals how skin and melanoma cells interact during melanoma spreading and invasion. J Theor Biol. 2017;423:13–25. doi: 10.1016/j.jtbi.2017.04.017

18. Simpson MJ, Towne C, McElwain DLS, Upton Z. Migration of breast cancer cells: understanding the roles of volume exclusion and cell-to-cell adhesion. Phys Rev E. 2010;82:041901. doi: 10.1103/PhysRevE.82.041901

19. Gerlee P. The model muddle: in search of tumor growth laws. Cancer Res. 2013;73:2407–2411. doi: 10.1158/0008-5472.CAN-12-4355

20. Vo BN, Drovandi CC, Pettitt AN, Simpson MJ. Quantifying uncertainty in parameter estimates for stochastic models of collective cell spreading using Approximate Bayesian Computation. Math Biosci. 2015;263:133–142. doi: 10.1016/j.mbs.2015.02.010

21. Green JEF, Waters SL, Whiteley JP, Edelstein-Keshet L, Shakesheff KM, Byrne HM. Non-local models for the formation of hepatocyte-stellate cell aggregates. J Theor Biol. 2010;267:106–120. doi: 10.1016/j.itbi.2010.08.013

22. Chan KS, Koh CG, Li HY. Mitosis-targeted anti-cancer therapies: where they stand. Cell Death Dis. 2012;3:e411. doi:10.1038/cddis.2012.148

23. Ramaraj P. *In vitro* inhibition of human melanoma (BLM) cell growth by progesterone receptor antagonist RU-486 (Mifprestone). J Caner Ther. 2016;7:1045–1058. doi: 10.4236/ict.2016.713101

24. Lai X, Friedman A. Combination therapy for melanoma with BRAF/MEK inhibitor and immune checkpoint inhibitor: a mathematical model. BMC Syst Biol. 2017;11:70. doi: 10.1186/s12918-017-0446-9

25. Dawson RA, Upton Z, Malda J, Harkin DG. Preparation of cultured skin for transplantation using insulin-like growth factor I in conjunction with insulin-like growth factor binding protein 5, epidermal growth factor, and vitronectin. Transplantation. 2006;81:1668–1676. doi: 10.1097/01.tp.0000226060.51572.89

